# Artificial intelligence virtual cell immune recovery model for screening traditional Chinese medicine ingredients

**DOI:** 10.64898/2026.06.16.732528

**Authors:** Chengpeng Hu, Beier Xiao, Calvin Yu-Chian Chen

## Abstract

Screening therapeutic candidates from single-cell transcriptomes requires a target that is closer to treatment response than disease-signature reversal. In immune diseases, post-treatment recovery may follow patient- and lineage-specific trajectories rather than a simple return along the pretreatment disease axis. We developed ImmuneNavi, an artificial intelligence virtual cell (AIVC) immune recovery model for ranking traditional Chinese medicine ingredients from paired PBMC data. The model maps heterogeneous PBMC cohorts to a common healthy immune coordinate system, constructs patient-lineage disease and recovery states, and processes ITCM treated-control profiles into a fixed ingredient perturbation bank. Patient and ingredient states are represented in matched gene, pathway and transcription-factor views, allowing the model to combine local transcriptional direction with more stable program-level features. A matcher trained on one paired treatment cohort preserved recovery-aligned ingredient rankings in independent PBMC cohorts without redefining the feature space, candidate set or preprocessing procedure. ImmuneNavi provides an AIVC model that uses paired immune-state measurements to screen natural-product candidates for experimental follow-up.

## 1 Introduction

Recent artificial intelligence virtual cell (AIVC) models are beginning to turn cellular atlases into systems for reasoning about intervention. These models can represent cellular states and predict responses to genetic, chemical or environmental perturbations, but screening therapeutic candidates adds a further requirement: the predicted or matched perturbation has to be judged against a treatment-relevant endpoint [1– 5]. This distinction matters in immune disease. A model may predict a change in cell state, yet still leave open whether that change resembles recovery in the patient and lineage of interest. Recent benchmarks show that extrapolation across biological contexts remains difficult, especially when disease background, treatment and platform all change [6].

Transcriptomic drug repurposing has usually defined that endpoint through signature opposition. Connectivity Map and LINCS established that disease and compound perturbation signatures can be compared at scale, and inverse drug-disease connectivity has become a practical way to nominate candidates [7, 8]. Single-cell extensions and perturbation models add cell-type resolution, patient-level heterogeneity or counterfactual response prediction [9–14]. These approaches are useful when reversing a disease signature is a suitable surrogate. They do not define which perturbation best matches the immune movement that is actually observed after treatment.

Inflammatory biology supports treating this as a separate target. Inflammation is an organized response to infection or tissue damage, and its resolution is not a passive decay of inflammatory activation. It involves active cellular and molecular programs that restore immune homeostasis [15–17]. In chronic inflammatory disease, defective resolution can sustain pathology after the initiating stimulus has changed [18]. A compound that opposes part of a pretreatment inflammatory profile may therefore differ from a compound whose perturbation profile resembles the regulatory or pro-resolving movement seen after therapy.

Paired pre- and post-treatment peripheral blood mononuclear cell (PBMC) data make this target measurable. For a given patient and immune cell type, the pretreatment state defines disease-associated deviation, whereas the post-minus-pre transition records the immune-state movement associated with treatment. A healthy reference places PBMC cohorts collected in different diseases and platforms into one coordinate system. Human reference atlases, atlas-level integration benchmarks and reference-mapping approaches provide normal cellular baselines for such comparisons [19–24]. The healthy immune reference defines a common space in which pretreatment deviation, reference-restoration and observed-recovery movement can be separated.

Traditional Chinese medicine (TCM) and natural products are a practical setting for this ranking problem. The candidate ingredient space is large, while experimental validation can test only a small fraction of plausible compounds. Network pharmacology has helped organize herbs, ingredients, targets and pathways into interpretable maps [25–27]. ITCM adds a complementary pharmacotranscriptomic resource by providing treated-control expression profiles for active TCM ingredients [28]. These resources identify candidate mechanisms and perturbation states, but they do not specify which ingredient should be tested first for a patient-lineage immune recovery trajectory.

Here we introduce ImmuneNavi, an AIVC immune recovery model for screening TCM ingredients in PBMC immune-state space. Each query is a patient-lineage pretreatment state. Each candidate is an ITCM ingredient perturbation state, defined as the treated-minus-control transcriptional response of an ingredient and represented in the same gene, pathway and transcription-factor views as the PBMC states. The ranking endpoint is agreement with the observed-recovery transition. We use paired PBMC cohorts to show that this endpoint differs from reference-restoration and disease-signature reversal, train ImmuneNavi to rank 496 ingredients, and test whether the learned ranking transfers to external PBMC datasets without cohort-specific retraining.

## 2 Results

### 2.1 State construction

To compare patient immune states with ingredient perturbation states, we first constructed a healthy-reference-anchored multi-view state space (Fig. 2). Paired PBMC profiles were mapped to the healthy immune reference and aggregated by patient, time point and immune lineage. For each patient-lineage query, this produced three related vectors: the pretreatment disease deviation, the reference-restoration direction towards the matched healthy centroid and the observed-recovery transition.

**Fig. 1.**
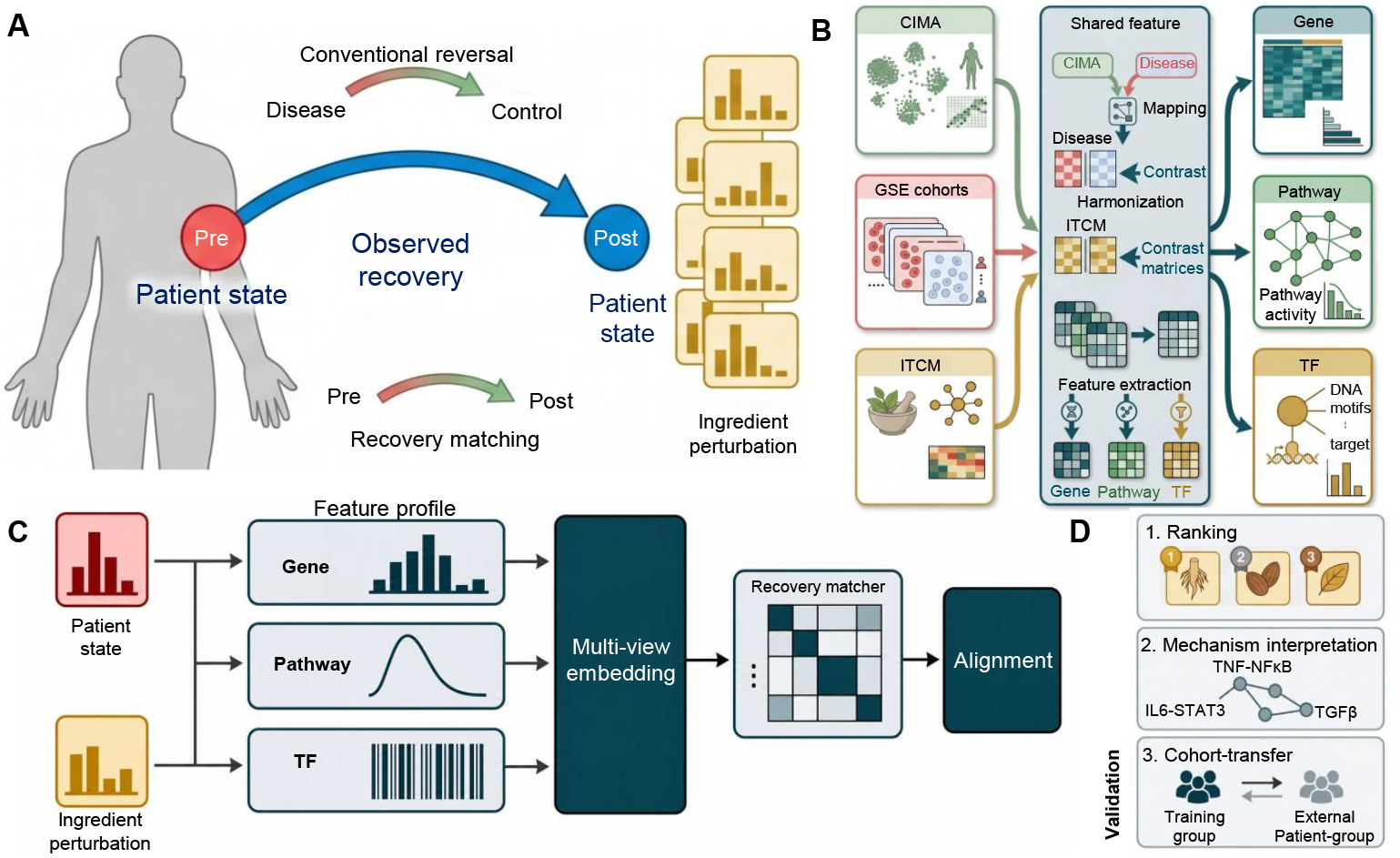
ImmuneNavi for healthy-reference-anchored multi-view recovery matching. **(A)** Paired single-cell disease cohorts are mapped to a healthy immune reference to define patient- and lineage-specific disease, reference-restoration and observed-recovery states. **(b)** ITCM treated-control perturbation profiles are processed into ingredient-level states and represented in the same feature space as patient states. **(c)** Gene, pathway and transcription-factor views provide complementary coordinates for comparing heterogeneous patient and perturbation data. **(d)** ImmuneNavi ranks candidate ingredients for each patient-lineage query and tests transfer across internal and external cohorts.

**Fig. 2.**
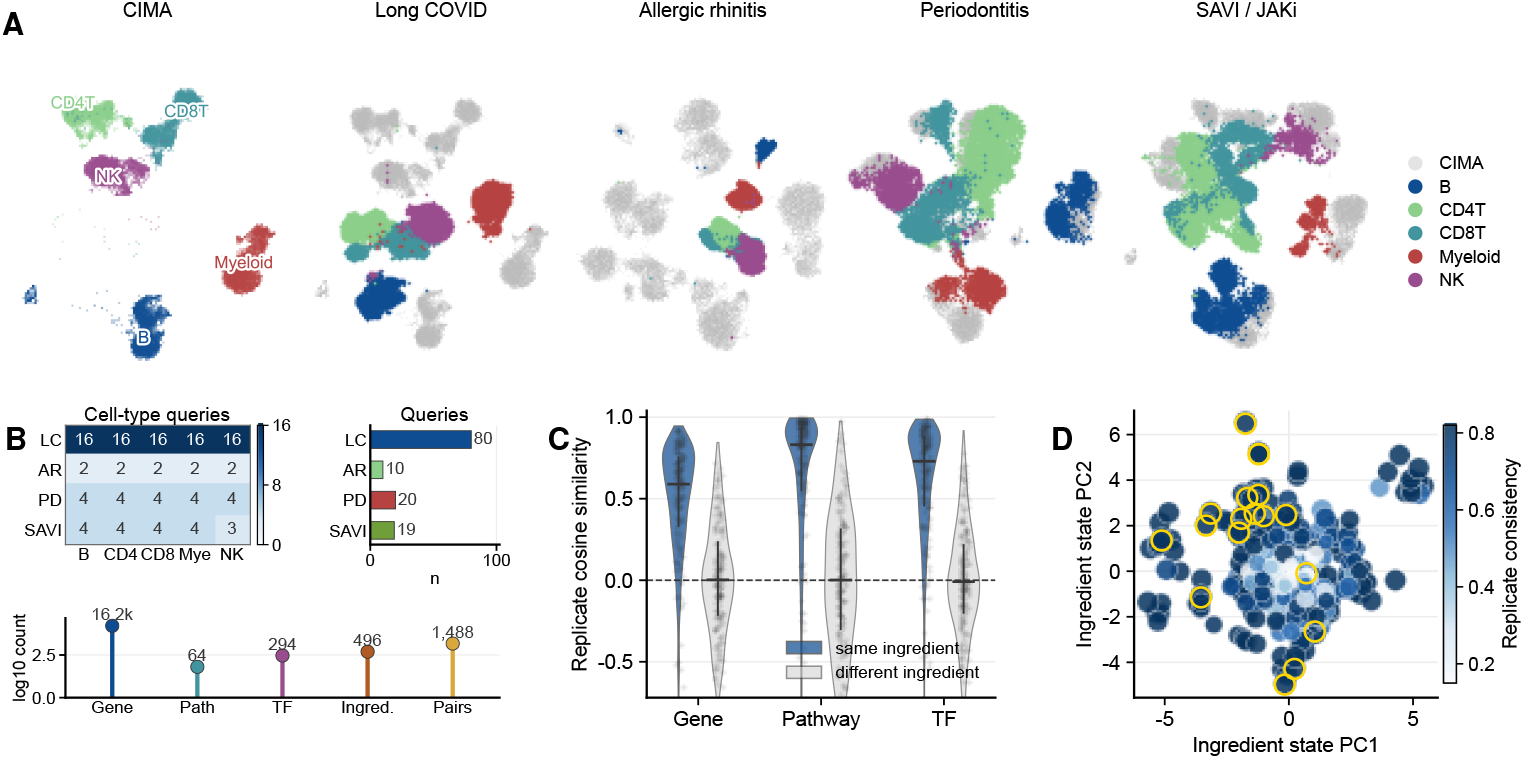
Shared state space for recovery queries and ITCM perturbations. **(A)** Reference-mapped PBMC profiles show the healthy atlas and four paired treatment cohorts: long COVID, allergic rhinitis, periodontitis and SAVI/JAK-inhibitor. **(B)** Query counts by cohort and cell lineage are shown with the frozen gene, pathway, transcription-factor and ITCM feature sets. **(C)** Same-ingredient perturbation pairs are more consistent than unrelated pairs across the three views. **(D)** The ITCM candidate bank spans a structured perturbation landscape, with reproducibility encoded by point color.

Five public single-cell datasets had distinct roles in this space. The COVID-19 immune atlas (GSE158055) was used for Stage-I pretraining. The long COVID herbal-therapy cohort (GSE265753) supplied the supervised recovery-matching task. The Peiyuan Tong-qiao decoction allergic-rhinitis cohort (GSE273975), the non-surgical periodontal-therapy periodontitis cohort (GSE174609) and the SAVI/JAK-inhibitor cohort (GSE226598) were held out for external transfer. The same five immune lineages were retained across the paired treatment cohorts. On the ingredient side, ITCM treated-control profiles were converted into perturbation-pair states and averaged into a 496-ingredient candidate bank. The final feature panel was frozen from the overlap among the COVID-19 atlas, the long COVID cohort and ITCM. This yielded a shared representation with gene, pathway and transcription-factor views. Reference-mapping quality, retained-lineage recall and external feature-projection checks are reported in Supplementary Fig. 1.

The three views captured different levels of the comparison. Gene-level states preserved high-resolution direction, whereas pathway and transcription-factor activities summarized coordinated signaling and regulatory variation across platforms and cell contexts. Stage-I pretraining used COVID-19 disease states and ITCM perturbation pairs to expose the encoder to both immune-state and ingredient-state variation before Stage-II recovery matching on the paired long COVID cohort.

### 2.2 Recovery matching

We next tested whether ITCM perturbation states could be ranked against the observed-recovery movement in this shared space. Endpoint controls separated observed recovery from two simpler targets: reference-restoration and reversal of the pretreatment disease deviation. Observed recovery was only partly aligned with reference-restoration, and top-ranked ingredients differed from those selected by disease-signature reversal. The healthy reference therefore provided the coordinate system, whereas the paired observed-recovery transition supplied the ranking target. Additional endpoint diagnostics quantifying observed recovery, reference-restoration and reversal-style rankings are provided in Supplementary Fig. 2.

With the 496-ingredient bank and the observed-recovery endpoint fixed, ImmuneNavi ranked ingredients more consistently with paired recovery than reversal-based and single-view alternatives (Fig. 3). Patient-averaged nDCG@10 was 0.755 in long COVID validation and 0.713, 0.732 and 0.660 in the allergic-rhinitis, periodontitis and SAVI/JAK-inhibitor transfer cohorts, respectively. This pattern was observed in patient-grouped validation within the long COVID cohort and in all three external PBMC cohorts without cohort-specific retraining. The external tests changed disease context and treatment setting: one cohort measured allergic rhinitis after Peiyuan Tong-qiao decoction, one measured periodontitis after non-surgical periodontal therapy, and one measured SAVI after JAK-inhibitor treatment. In all cases, post-treatment profiles were used only to compute evaluation labels.

**Fig. 3.**
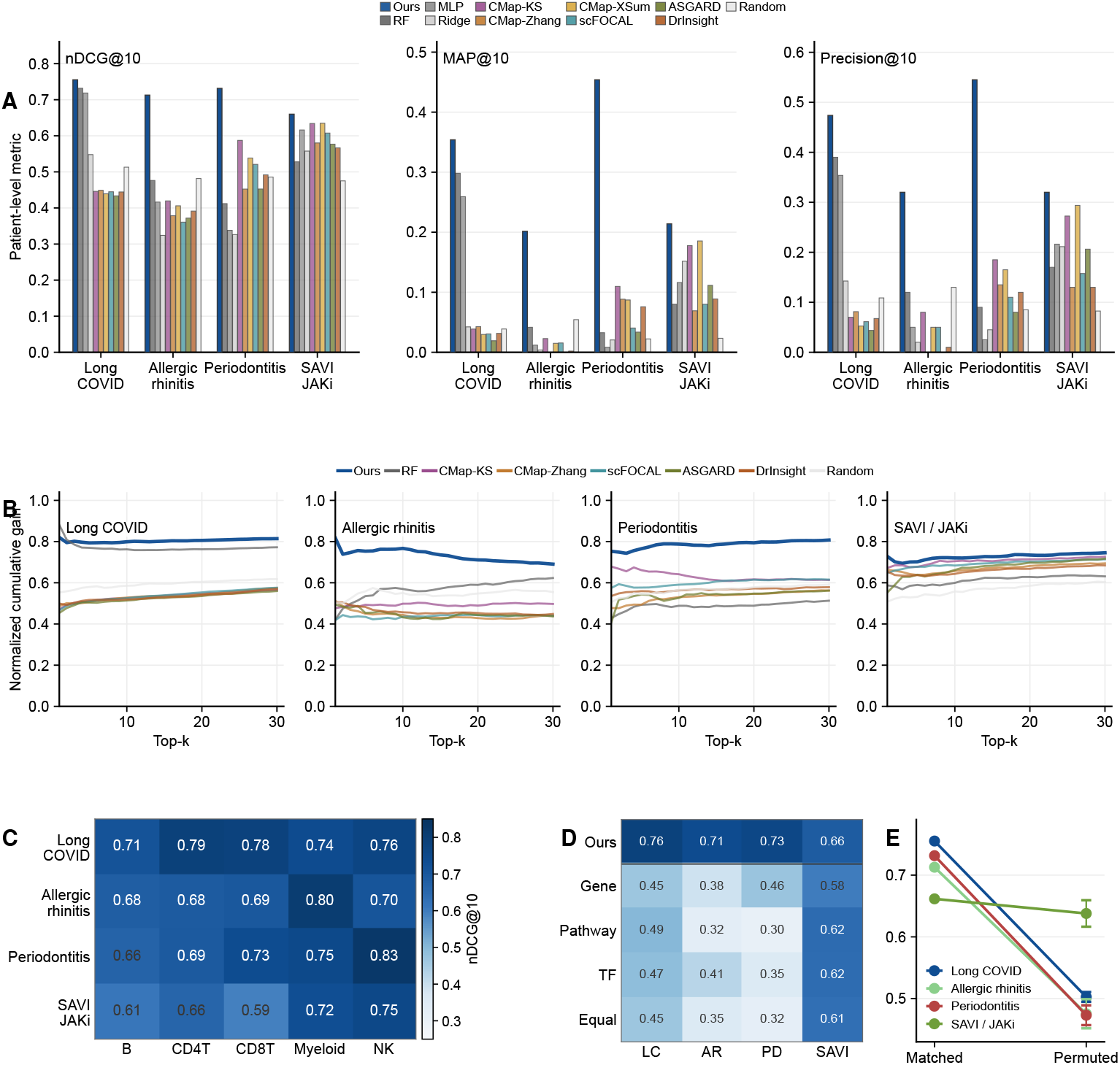
Recovery-aligned ingredient ranking across four PBMC cohorts. **(A)** Patient-level nDCG@10, MAP@10 and precision@10 compare ImmuneNavi with baseline methods. **(B)** Top-k gain curves show enrichment of observed-recovery-relevant ingredients across cohorts. **(C)** ImmuneNavi performance by immune lineage. **(D)** Multi-view ranking compared with single-view and equal-fusion ablations. **(E)** Matched observed-recovery labels compared with permuted labels.

Fig. 3 also tested whether this transfer depended on a single lineage or view. Ranking performance was retained across the five immune lineages. Gene-only, pathway-only, transcription-factor-only and equal-fusion variants showed lower recovery ranking than ImmuneNavi in the internal and external cohorts. Label permutation reduced the scores towards the random-ranking range. These controls indicate that the external rankings depended on a healthy-reference-anchored multi-view recovery signal rather than a dataset-specific shortcut or a generic ranking bias. Patient-level uncertainty, secondary ranking metrics, lineage-specific performance, endpoint-weight sensitivity and pretraining-related dataset-shift analyses are provided in Supplementary Figs. 3 and 4.

### 2.3 Biological analysis

We then examined ImmuneNavi rankings at patient-lineage resolution (Fig. 4). In the long COVID cohort, ingredients with high mean model scores included Paeonol, Albiflorin, Pulegone, Picroside II, Pseudoephedrine, Salicylic acid, Muscone and Harpagoside. Scores varied across immune lineages and patients, and the model did not reduce the cohort to a single ingredient list.

**Fig. 4.**
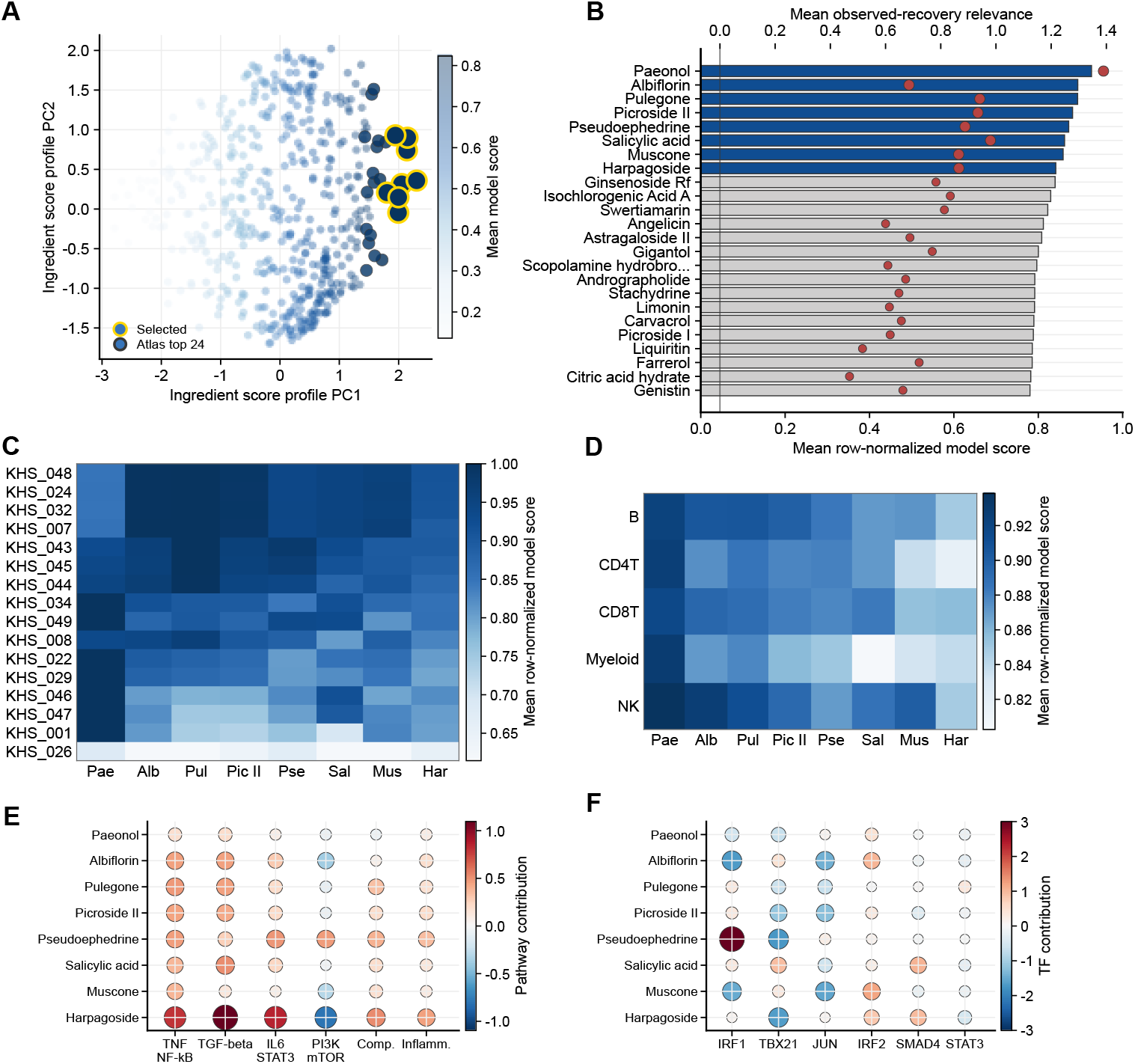
Ingredient rankings resolve cohort-level candidates, patient variation and recovery-program annotations. **(A)** The ingredient atlas summarizes candidate score profiles across recovery queries. **(B)** Leading candidates are compared with observed-recovery relevance. **(C)** Patient-level profiles show how candidate rankings vary across individuals. **(D)** Cell-lineage profiles show how nominated ingredients vary across immune contexts. **(E)** Pathway summaries annotate recovery programs associated with selected perturbations. **(F)** Transcription-factor summaries provide complementary regulatory annotations.

Representative myeloid queries provided query-level examples of this ranking behavior. In the long COVID cohort, several top-ranked ingredients also ranked highly under direct observed-recovery relevance but poorly under disease-reversal relevance. These examples show how ranked candidate perturbations arise from individual patient-lineage states and can be taken forward for follow-up testing.

Feature-level summaries annotated these nominations with immune programs. Signed pathway contributions measured whether a candidate perturbation and the observed recovery transition moved in the same direction along pathway features. The selected directional pathway panel included TNF-*α* signaling via NF-*κ*B, IL6–JAK– STAT3 signaling, TGF-*β* signaling and inflammatory response. The transcription-factor panel, including TBX21, IRF1, JUN, SMAD4, IRF2 and STAT3, was used as annotation rather than enrichment evidence. Many leading ImmuneNavi nominations had prior anti-inflammatory or immunomodulatory support; representative examples include Paeonol and andrographolide, which have been linked to inflammatory or immune-regulatory pathways in pharmacological reviews [29, 30]. Literature support alone did not determine the ranking: curcumin, a well-studied immunomodulatory natural product, was ranked lower, as expected, because ImmuneNavi screens candidates by their agreement with patient- and lineage-specific observed recovery rather than generic anti-inflammatory activity [31]. Formula-level hit rate provided an additional check against curated TCM ingredient lists, but it behaved differently across cohorts. We therefore treated it as auxiliary evidence rather than as the primary endpoint. Formula-level retrieval checks and additional evidence-layer annotations for the internal cohort are shown in Supplementary Fig. 5.

## 3 Discussion

This study presents ImmuneNavi as an AIVC immune recovery model that frames TCM ingredient screening as a recovery-aligned state-matching problem in PBMC immune-state space. Candidate ingredients were ranked by the concordance between ITCM-derived perturbation states and the observed-recovery movement of each patient-lineage query, with reversal-based retrieval retained as a baseline for comparison [7, 8]. The resulting ranking asks whether a candidate perturbation resembles the immune transition observed after treatment, a question consistent with inflammatory resolution being an active and regulated process rather than a simple negation of pretreatment inflammation [16–18].

A reproducible workflow for placing new paired datasets in the same immune-state space is therefore a necessary part of the work [32]. Healthy single-cell references provide the basis for comparing disease states across studies [21, 23, 24]; here, the reference layer standardizes state construction before ingredient ranking. The same reference mapping, feature construction and ITCM candidate bank define the query state, reference-restoration direction and observed-recovery direction across cohorts. A new paired PBMC dataset can therefore be processed through the same healthy-reference-anchored workflow without redefining the state space or candidate set for each study. Future cohorts can contribute new recovery labels while reusing the same preprocessing and candidate space across different disease settings.

The ImmuneNavi matcher adds a second layer of generalization to this workflow. Once patient and ingredient states are represented in the same multi-view space, the model learns how gene-level direction, pathway activity and transcription-factor activity jointly relate to recovery-aligned ranking. This three-view representation helps transfer across cell types and datasets: gene features retain local expression direction, whereas pathway and transcription-factor views summarize higher-level immune programs. It also reduces dependence on a single feature scale when cell context and platform effects vary between PBMC datasets. The external validation experiments test this deployment pattern directly: a model trained on long COVID recovery was applied to allergic-rhinitis, periodontitis and SAVI/JAK-inhibitor PBMC cohorts without cohort-specific retraining. Because the candidate bank, feature views and ranking definitions were held fixed, these experiments evaluate transfer across paired PBMC studies rather than a new manual screening exercise for each cohort. The pretraining analysis in Supplementary Fig. 4 further separates frozen external transfer from dataset-shift effects between the long COVID training manifold and the external cohorts. This places the approach alongside single-cell drug-repurposing and perturbation-response methods [12–14], while extending the use case to a fixed natural-product perturbation bank and paired patient recovery trajectories.

For TCM and natural-product studies, this combination of a common processing workflow and a transferable matcher provides a way to move from broad candidate catalogs to patient-state-guided screening. Network pharmacology and pharmaco-transcriptomic resources have organized herbs, ingredients, targets and perturbation profiles into computable knowledge bases [25–28]. The present framework adds a recovery-state filter to that information by asking which perturbation profiles are compatible with measured immune-state improvement in paired data. In practice, this filter can help decide which ingredients merit functional testing first, especially when a curated resource contains many plausible candidates with overlapping target or pathway annotations.

The current results should be interpreted as transcriptomic screening results. They support recovery-aligned matching as a strategy for nominating ingredients, but they do not establish clinical efficacy or causal mechanism. The next step is to connect this ranking framework to prospective paired cohorts with clinical response measurements, functional immune-cell perturbation assays, perturbation resources with explicit dose and time structure, and models that represent formula combinations rather than single ingredients alone. These extensions would test whether a healthy-reference-anchored recovery-matching workflow can improve the selection of natural-product candidates in settings where patient immune states, perturbation profiles and treatment outcomes are measured together.

## 4 Methods

### 4.1 Data resources and task definition

Candidate ingredients were ranked for each patient-lineage query. A query (*i, c*) was defined by a pretreatment PBMC state and immune lineage, and the candidate set *A* comprised ITCM ingredient perturbation states. ImmuneNavi received the healthy-reference-anchored disease state for the query and each candidate ingredient perturbation state, then returned a score *s*_*i,c,a*_ for every ingredient *a* ∈ *A*. Observed pre-to-post recovery vectors defined the supervised target and primary evaluation endpoint; reference-restoration directions defined a secondary reference endpoint. At prediction time, rankings were produced from the pretreatment query state and candidate ingredient states.

The Chinese Immune Multi-Omics Atlas (CIMA) was used as the healthy immune reference for lineage-specific baselines [24]. Disease and paired pre/post immune states were assembled from five public single-cell studies. A COVID-19 immune atlas was used for representation pretraining (GEO: GSE158055) [33]. A paired peripheral-blood study of long COVID patients receiving herbal therapy was used for supervised recovery matching (GEO: GSE265753) [34]. External validation used an allergic-rhinitis PBMC study of Peiyuan Tong-qiao decoction (GEO: GSE273975) [35], a periodontitis PBMC study sampled before and after non-surgical periodontal therapy (GEO: GSE174609) [36], and a SAVI PBMC study sampled before and after JAK-inhibitor treatment (GEO: GSE226598) [37]. Candidate perturbation states were obtained from ITCM, which contains pharmacotranscriptomic profiles for active TCM ingredients [28].

### 4.2 Healthy-reference-anchored immune states

Query cells were mapped to the healthy immune reference with Scanpy preprocessing and scANVI reference mapping [38, 39]. Five circulating immune lineages were retained: B cells, CD4 T cells, CD8 T cells, myeloid cells and NK cells. Cells were aggregated into pseudobulk profiles by patient, time point and cell type. Each pseudobulk state was represented in three views, *v* ∈ *V* = {*g, p, t*} , corresponding to signed gene ranks, pathway activities and transcription-factor activities. This multi-view representation retained within-profile gene-level direction while summarizing coordinated signaling and regulatory changes in lower-dimensional pathway and transcription-factor spaces. Pathway activities were inferred from PROGENy signaling footprints and MSigDB Hallmark gene sets, and transcription-factor activities were inferred from DoRothEA regulons using decoupleR [40–43].

Let 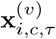 denote the pseudobulk profile of patient *i*, cell type *c*, time point *τ* and view *v*. Let 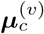 denote the mean healthy-reference pseudobulk state for cell type *c* and view *v*. The disease deviation state and the two recovery endpoints were defined as

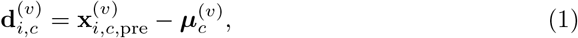

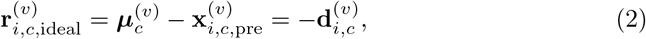

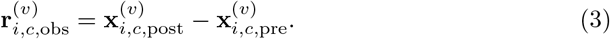

Expressing both time points as deviations from the matched healthy centroid gives the same observed vector because the same cell-type centroid is used for the pair. The reference-restoration endpoint measured the direction from the pretreatment state back to the matched healthy centroid. The supervised training set contained 80 observed-recovery queries from 16 long COVID patients. External validation contained 10 paired patient-lineage queries from the allergic-rhinitis cohort, 20 from the periodontitis cohort and 19 from the SAVI/JAK-inhibitor cohort.

### 4.3 Ingredient perturbation states

The ITCM treated-control expression profiles were converted into perturbation states in the same three-view space as the patient states. For an ingredient *a* with replicate treated-control pairs R_*a*_, the ingredient state was

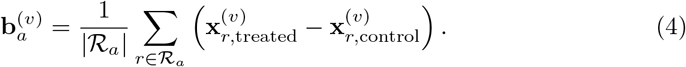

For the gene view, treated-control deltas were converted to signed within-profile ranks so that patient and ingredient vectors were compared by direction rather than absolute expression scale. Pathway and transcription-factor views were retained as activity differences. The frozen feature panel was the intersection shared by the COVID-19 pretraining states, the long COVID recovery states and the ITCM perturbation states, comprising 16,229 genes, 64 pathway features and 294 transcription-factor features. This produced 1,488 ITCM pair states and 496 ingredient-level candidate states.

### 4.4 Recovery-alignment relevance score

For supervised learning and evaluation, the observed-recovery relevance of ingredient *a* for query (*i, c*) was defined by cosine agreement between the ingredient perturbation state and the observed recovery vector across the three views. For each view,

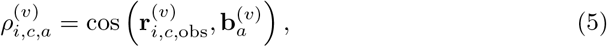

and the multi-view relevance score was

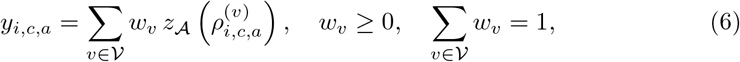

where *z*_*A*_(·) denotes standardisation across all candidate ingredients for the same query. The soft training target was

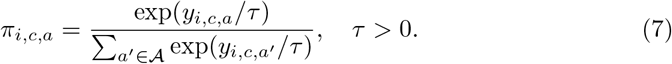

The secondary reference-restoration endpoint used the same equations after replacing 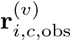 with 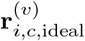. The weight vector *w* and temperature *τ* were fixed before model fitting and were not learned from the validation or external cohorts.

### 4.5 ImmuneNavi model

ImmuneNavi consisted of a multi-view state encoder, a cell-type-conditioned recovery generator and a cross-attention ingredient matcher. For each view, the input vector was divided into fixed-length chunks and passed through a Transformer encoder [44]. The view embeddings were then fused into a shared latent state:

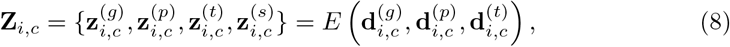

where *s* denotes the shared fused state. A residual generator *G*, conditioned on a learned cell-type embedding **e**_*c*_, predicted latent recovery tokens:

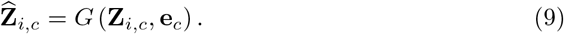

Ingredient states were encoded with the same encoder to form **Z**_*a*_ = *E*(**b**_*a*_). The matcher used bidirectional cross-attention between 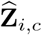 and **Z**_*a*_, followed by pooled attended features, element-wise products, absolute differences and a cosine term, to produce the predicted ingredient score *s*_*i,c,a*_. Candidate ingredients were ranked in descending order of *s*_*i,c,a*_.

### 4.6 Training procedure

Training was performed in two stages. Stage I pretrained the encoder on 902 healthy-reference-anchored disease states from the COVID-19 immune atlas and 1,488 ITCM pair states. The pretraining objective combined masked-input reconstruction, cross-view alignment, contrastive consistency between two stochastic masked views and reference-restoration pairing:

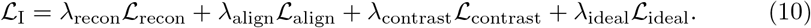

ℒ_ideal_ matched the latent state predicted from a disease state to the encoded reference-restoration state. Stage II trained the recovery generator and matcher on the 80 observed-recovery queries. For each query, the predicted scores were compared with the soft relevance target *π* by cross-entropy:

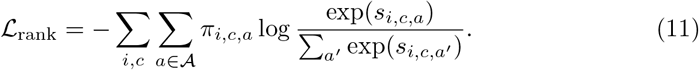

The ranking loss was combined with a latent target-matching loss and, for the pretrained model, an anchor loss:

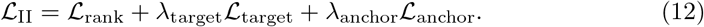

ℒ_target_ was the average cosine distance between generated latent recovery tokens and encoded observed-recovery tokens for the shared, pathway and transcription-factor views. ℒ_anchor_ was the mean squared deviation between the fine-tuned encoder and the frozen Stage I encoder for the selected anchored states. Patient-grouped folds were used throughout supervised training, so all cell-type queries from a patient were assigned to the same fold. The model used 128-dimensional latent tokens, gene chunks of 32 features, pathway and transcription-factor chunks of 8 features, two view-encoder layers and two fusion layers.

### 4.7 Evaluation and baseline comparisons

The primary endpoint was patient-averaged nDCG@10 against the observed recovery relevance score. For a query, relevance values were shifted and rescaled to [0, 1], and

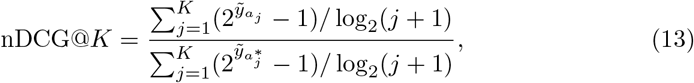

where *a*_*j*_ is the ingredient ranked at position *j* by the model and 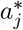 is the ingredient at position *j* in the oracle ordering induced by 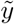. Query-level metrics were averaged within each patient and then averaged across patients with equal patient weights. Secondary endpoints included nDCG@20, top-10 observed-recovery alignment, Spearman correlation and the same metrics against the reference-restoration endpoint.

External validation applied the frozen fold ensemble trained on long COVID recovery to the allergic-rhinitis, periodontitis and SAVI/JAK-inhibitor cohorts without cohort-specific retraining. Comparator rankings included random and shuffled controls, disease-reversal scores derived from CMap/LINCS connectivity logic [7, 8], supervised regression baselines and view-ablation variants. The disease-reversal comparators used − 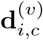 as the target vector. All comparators used the same 496-ingredient ITCM candidate bank and were evaluated with the same observed-recovery and reference-restoration relevance definitions. Confidence intervals were estimated by 1,000 patient bootstraps, and empirical *P* values were calculated from 200 label-permutation runs.

## Declarations

### Data availability

All datasets used in this study are publicly available. The single-cell datasets were obtained from the Gene Expression Omnibus under accession codes GSE158055, GSE265753, GSE273975, GSE174609 and GSE226598. The healthy immune reference was derived from the Chinese Immune Multi-Omics Atlas, and candidate ingredient perturbation profiles were obtained from the public ITCM pharmacotranscriptomic resource. Source data supporting the figures are provided with this manuscript or can be regenerated from the cited public resources.

### Code availability

The code used for data processing, model training and external validation is available at GitHub: https://github.com/Youzhuqinghuan/ImmuneNav.

### Author contributions

C.H. conceived the study, designed and implemented the method, performed the main analyses, and wrote the manuscript. B.X. assisted with data curation, experimental analysis and validation, and preparation of the Supplementary Information. C.Y.-C.C. supervised the study and provided academic guidance and project support. All authors reviewed and approved the manuscript.

## Acknowledgements

The authors thank Zhida Chen and Chengen Jiang for helpful discussions.

