## Supplementary Information for "Artificial intelligence virtual cell immune recovery model for screening traditional Chinese medicine ingredients"

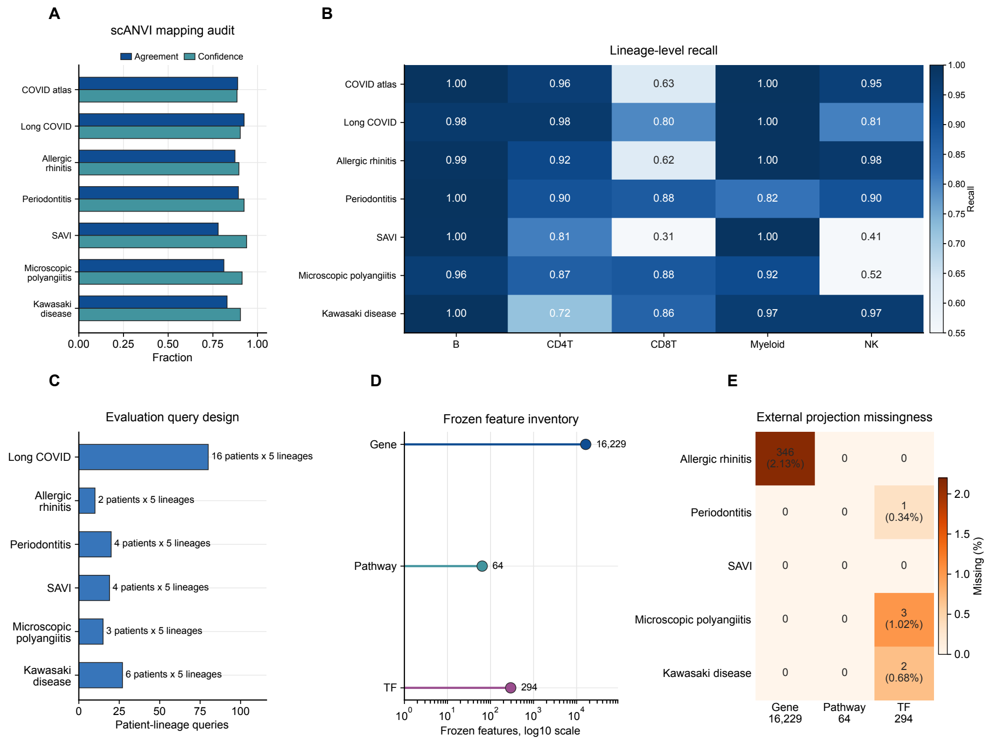


**Fig. 1 Reference mapping and fixed feature projection across disease cohorts.** All cohorts were mapped to a common healthy reference and represented with the same gene, pathway and transcription-factor features before recovery directions and ingredient perturbation states were compared. This figure reports the mapping, query-design and feature-projection checks used for that fixed state space. **A**, Cohort-level scANVI mapping audit for the COVID atlas and six disease cohorts. Bars show agreement between manual major-lineage annotations and transferred labels, together with median transferred-label confidence. **B**, Manual-lineage recall for the retained B, CD4 T, CD8 T, myeloid and NK compartments after reference mapping. **C**, Evaluation-query design for the internal long COVID cohort and five external cohorts. Bars show patient-lineage query counts; text gives patients per retained lineage and the number of retained lineages. **D**, Number of frozen gene, pathway and transcription-factor features used for each patient and ingredient vector. The x axis is log-scaled because the three views differ greatly in size. **E**, Feature missingness after each external cohort was projected onto the frozen feature order. Rows show the five external evaluation cohorts; columns show the gene, pathway and transcription-factor views.

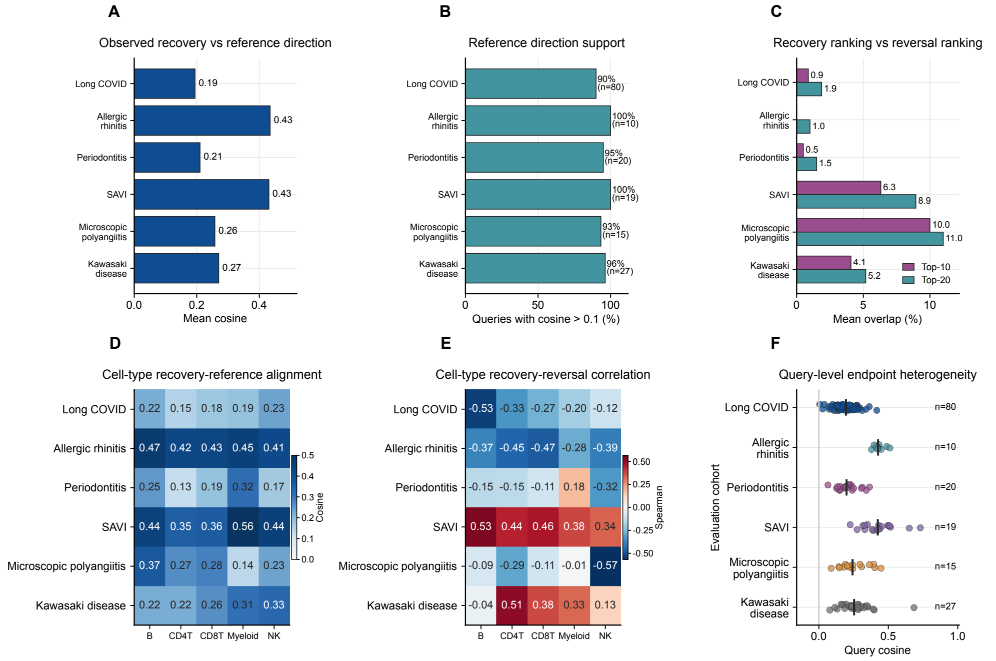


**Fig. 2 Measured paired recovery is distinct from direct healthy-reference or disease reversal.** The benchmark ranks ingredients by agreement with the observed post-minus-pre treatment response in each patient-lineage query. This figure compares that endpoint with two related quantities: the direction from pretreatment state to the healthy reference, and a disease-reversal ranking rule. These comparisons define how the reference is used, without replacing measured recovery as the ranking endpoint. **A**, Mean cosine similarity between observed paired recovery and the direct pretreatment-to-reference direction. Values below 1 indicate that measured recovery is not identical to direct reference restoration. **B**, Fraction of patient-lineage queries whose observed recovery vector points in the reference direction, defined as cosine $> 0.1$. This panel tests whether the reference provides a consistent directional anchor across cohorts. **C**, Mean Top-10 and Top-20 overlap between ingredient rankings induced by the observed-recovery endpoint and by a reversal-style endpoint. Low overlap indicates that the two endpoints rank different ingredients near the top. **D**, Cell-type-specific observed-versus-reference cosine values for the retained immune lineages. **E**, Cell-type-specific Spearman correlation between observed-recovery and reversal-style ingredient relevance. Weak or negative values indicate different ranking structure within the same lineage. **F**, Query-level observed-versus-reference cosine values. Each point represents one patient-lineage query, and vertical black ticks mark cohort medians. These panels support using measured paired recovery as the ranking endpoint, with reference and reversal analyses used as diagnostics.

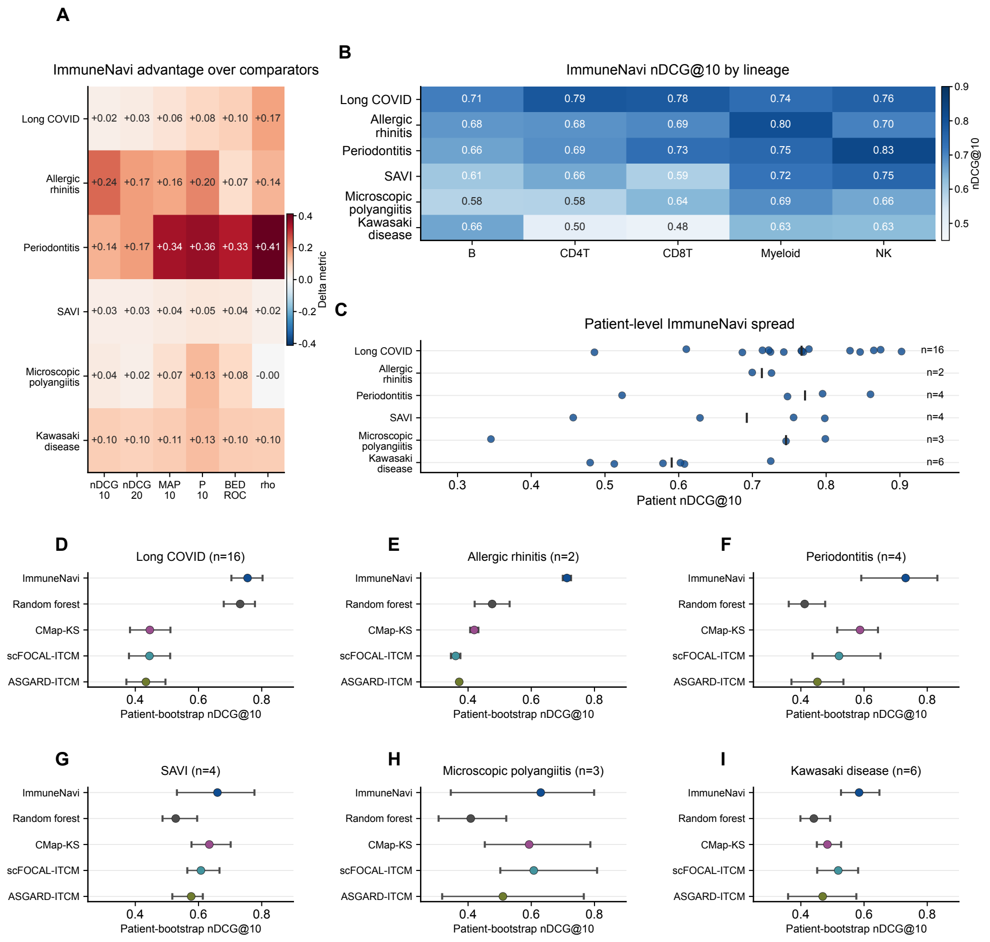


**Fig. 3** **Supplementary performance checks across metrics, lineages, patients and bootstrap uncertainty.** The main text reports the compact recovery-ranking comparison. This figure extends that comparison across the six evaluation datasets and shows where the signal appears across metrics, immune lineages and patients. **A**, Difference between the ImmuneNavi primary method and the better of random forest and CMap-KS for each ranking metric. Positive values indicate higher ImmuneNavi performance. **B**, ImmuneNavi nDCG@10 by retained immune lineage. **C**, Patient-level ImmuneNavi nDCG@10 values. Each point represents one patient, and vertical black ticks mark cohort medians. **D**–**I**, Dataset-specific patient-bootstrap nDCG@10 estimates for ImmuneNavi and four representative baselines. Points show bootstrap means, horizontal bars show confidence intervals, and panel titles give the cohort and patient count.

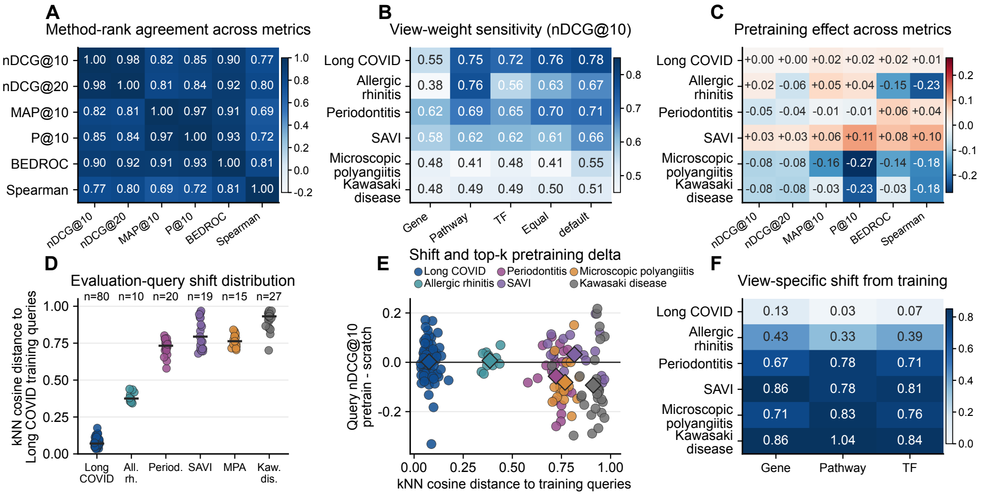


**Fig. 4** **Metric sensitivity and dataset heterogeneity of pretrained transfer.** **A**, Spearman correlations between method rankings induced by six recovery-ranking metrics. This panel tests whether the primary nDCG@10 endpoint is aligned with related top-k and correlation-based summaries. **B**, ImmuneNavi nDCG@10 under gene-only, pathway-only, TF-only, equal-weight and default multi-view endpoint definitions. This panel tests whether performance depends on one feature view or remains stable after gene, pathway and transcription-factor evidence are reweighted. **C**, Pretrained-minus-scratch deltas across datasets and metrics. Positive values favor the pretrained model, whereas negative values favor the scratch model. The pretrained model is the default ImmuneNavi setting and is favored in the internal long COVID cohort and selected external settings, whereas scratch can be higher in shifted cohorts or metrics. **D**, Query-level k-nearest-neighbor cosine distance from each evaluation query to the long COVID training-query manifold in the frozen multi-view state space; horizontal ticks mark medians. This panel measures dataset shift between the pretraining source and each evaluation cohort. **E**, Query-level relationship between the training-manifold distance in D and the pretrained-minus-scratch nDCG@10 delta in C; diamond markers indicate cohort means. This panel shows how reduced pretrained advantage is concentrated among more shifted queries and cohorts. **F**, Gene, pathway and TF components of the same training-manifold distance. Several added external cohorts show larger view-specific shifts than the long COVID cohort, especially in pathway and transcription-factor space. Both pretrained and scratch variants are frozen before external scoring, so scratch-favored cases are interpreted as context-dependent transfer under dataset heterogeneity, not as external-cohort tuning or leakage.

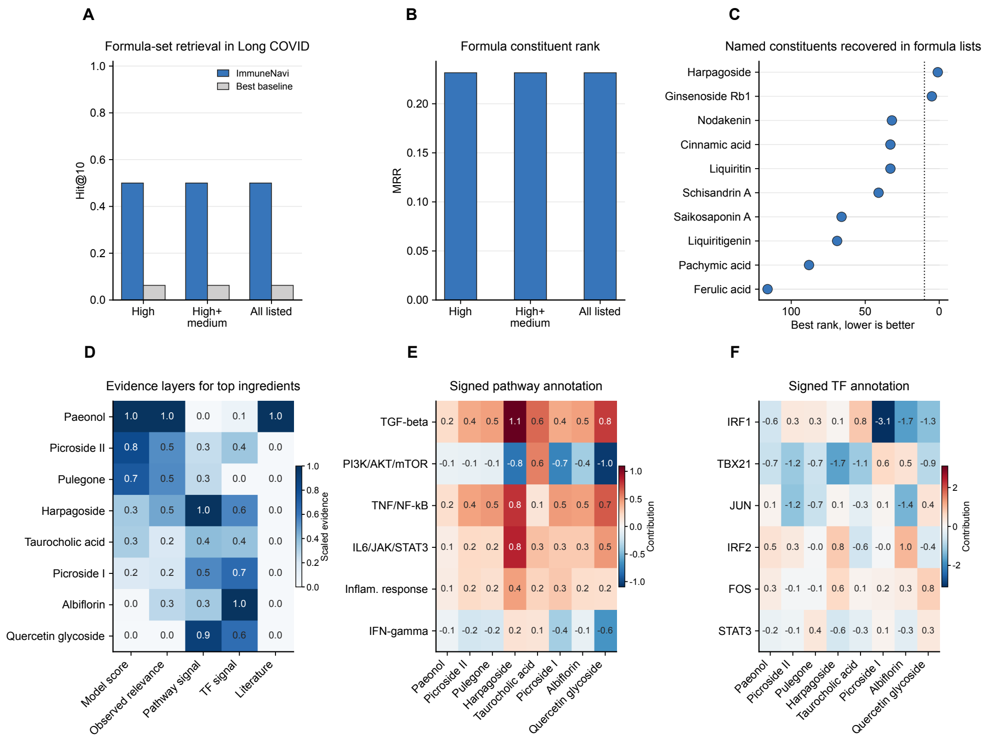


**Fig. 5** **Internal formula recovery and biological annotation of top-ranked ingredients.** **A**, Formula-level Hit@10 in the main long COVID experimental cohort. Curated treatment formula constituents were grouped into high-confidence, high-plus-medium and all-listed truth sets, and the bars test whether known formula constituents appear in the first ten ranked ingredients more often for ImmuneNavi than for the best-performing baseline in this analysis. **B**, Mean reciprocal rank for the same formula-truth definitions. This panel complements A by asking how early the first matching formula constituent appears, rather than only whether any match falls inside the top ten. **C**, Best ranks of named high-plus-medium formula constituents in the internal cohort. The dotted vertical line marks rank 10, matching the Hit@10 cutoff used in A. Ingredient names are shortened for readability, with full names retained in the source data. **D**, Evidence layers for the top eight internal ImmuneNavi-ranked ingredients. Model score, observed-recovery relevance, pathway contribution, transcription-factor contribution and curated literature support are scaled only for visual comparison within this panel; literature support is marked only where present in the curated source table. **E**, Signed pathway contributions for the same top ingredients across selected immune and inflammatory programs. **F**, Signed transcription-factor contributions for the same top ingredients. Panels D–F provide biological annotation of nominated ingredients, but they do not constitute causal mechanism validation; hit-rate analyses are restricted to the main experimental cohort.

### Supplementary Notes

#### Data processing and reference mapping

The Chinese Immune Multi-Omics Atlas (CIMA) was used as the healthy immune reference for mapping and lineage-specific baseline construction [1]. Query cells were mapped to this reference with a scVI/scANVI workflow implemented separately for each cohort [2, 3]. CIMA major-lineage labels were used as the supervised scANVI label field, whereas query cells were introduced as unknown-label cells during query adaptation. The mapped labels and confidence summaries were then used to aggregate patient-time-lineage pseudobulk states. This reference mapping creates a shared coordinate system for downstream recovery vectors, but it is not assumed to erase platform, disease or intervention heterogeneity.

#### Endpoint definitions

For a query $q = (i, c)$, let $\mathbf{x}^{(v)}_{\mathrm{pre},q}$ and $\mathbf{x}^{(v)}_{\mathrm{post},q}$ denote the pre- and post-treatment pseudobulk state in view $v$, and let $\boldsymbol{\mu}^{(v)}_c$ denote the CIMA healthy centroid for cell type $c$. The observed recovery direction is

$$\Delta^{(v)}_{\mathrm{obs},q} = \mathbf{x}^{(v)}_{\mathrm{post},q} - \mathbf{x}^{(v)}_{\mathrm{pre},q}.$$

The CIMA-ideal direction is

$$\Delta^{(v)}_{\mathrm{ideal},q} = \boldsymbol{\mu}^{(v)}_c - \mathbf{x}^{(v)}_{\mathrm{pre},q}.$$

The primary benchmark uses the observed recovery direction because it reflects the measured paired change. CIMA-ideal and disease-reversal analyses are reported as diagnostics because they test different biological assumptions.

#### Pretraining, dataset heterogeneity and dataset merging

Pretraining is the default ImmuneNavi setting and acts as a context-dependent regularizer. The internal long COVID cohort shows a modest pretrained advantage across several metrics, and selected external or correlation-style settings also favor the pretrained model. External transfer, however, is not uniform. Across the external cohorts, scratch can be higher on some top-k metrics when evaluation queries are farther from the long COVID training manifold. We interpret this pattern through dataset heterogeneity and transfer difficulty: disease background, intervention type, cohort size, cell-state distribution and feature projection all differ from the training setting. Because both variants are frozen before external scoring, this comparison does not introduce external-cohort training leakage. This is a dataset-merging risk in virtual-cell perturbation benchmarks: negative transfer can occur when perturbation context and biological state are not well aligned [4, 5].
